# Divergent genome evolution caused by regional variation in DNA gain and loss between human and mouse

**DOI:** 10.1101/179200

**Authors:** Reuben M Buckley, R Daniel Kortschak, David L Adelson

## Abstract

The forces driving the accumulation and removal of non-coding DNA and ultimately the evolution of genome size in complex organisms are intimately linked to genome structure and organisation. Our analysis provides a novel method for capturing the regional variation of lineage-specific DNA gain and loss events in their respective genomic contexts. To further understand this connection we used comparative genomics to identify genome-wide individual DNA gain and loss events in the human and mouse genomes. Focusing on the distribution of DNA gains and losses, relationships to important structural features and potential impact on biological processes, we found that in autosomes, DNA gains and losses both followed separate lineage-specific accumulation patterns. However, in both species chromosome X was particularly enriched for DNA gain, consistent with its high L1 retrotransposon content required for X inactivation. We found that DNA loss was associated with gene-rich open chromatin regions and DNA gain events with gene-poor closed chromatin regions. Additionally, we found that DNA loss events tended to be smaller than DNA gain events suggesting that they were more tolerated in open chromatin regions. GO term enrichment in human gain hotspots showed terms related to cell cycle/metabolism, human loss hotspots were enriched for terms related to gene silencing, and mouse gain hotspots were enriched for terms related to transcription regulation. Interestingly, mouse loss hotspots were strongly enriched for terms related to developmental processes, suggesting that DNA loss in mouse is associated with phenotypic changes in mouse morphology. This is consistent with a model in which DNA gain and loss results in turnover or “churning” of regulatory regions that are then subjected to selection, resulting in the differences we now observe, both genomic and phenotypic/morphological.

## Introduction

Evolution as a result of natural selection has led to many streamlined forms which follow directly from their function. However, in the case of genome evolution of complex organisms this connection is not quite so direct. One example is the evolution of genome size. In vertebrates, gene content has remained relatively constant, while the fraction of non-coding DNA varies drastically (Gregory 2005; Elliott and Gregory 2015; Gregory 2001). This observation is at the heart of the C-value enigma and raises many questions regarding the molecular drivers and evolutionary impacts of genome size variation. The major factor contributing to the total non-coding DNA genomic fraction is transposon load, due to mobile DNA elements that have actively replicated throughout evolution (Gregory 2001; Elliott and Gregory 2015). In humans, since their divergence from the common placental ancestor, transposon activity has caused approximately 815 Mb of DNA gain, almost one third of their extant genome (Kapusta et al. 2017; Lander et al. 2001). However, this is not the only factor driving genome size evolution. DNA loss via deletion also plays a role, with approximately 650 Mb of the human genome being lost over the same time period (Kapusta et al. 2017). Across mammals and birds these two forces operate in opposition to each other leading to the accordion model of genome evolution, where departures from this DNA gain and loss equilibrium cause genomes to either grow or shrink (Kapusta et al. 2017). Importantly, our understanding of DNA gain and loss stems from genome-wide estimates rather than detection of individual events. Therefore, the role of genome structure on widespread DNA gain and loss and its subsequent impact on lineage-specific species evolution remains unknown.

The ‘accordion’ model of genome size evolution raises important questions regarding the roles of natural selection and genetic drift. Genome size, like any other heritable trait, is shaped by a combination of both of these factors (Lynch and Walsh 2007). However, the contribution of each mechanism in diverse taxa remains an open question in biology, with evidence to support the impact of each (Whitney and Garland Jr 2010). For genome evolution driven by selection there are observations of various phenotypic correlates consistent across both mammals and birds. One example is the evolution of powered flight in bats and birds which requires a high metabolic rate. Because metabolism is more efficient in smaller cells, it has been suggested that in flying species there is particularly strong selection pressure against genome growth (Wright et al. 2014; Vinogradov and Anatskaya 2006; Kapusta et al. 2017). Alternatively, observed genome size variation can result from neutral evolutionary processes. Many higher order vertebrates have low effective population sizes resulting from reduced efficiency of selection (Lynch and Conery 2003), suggesting that neutral or mildly deleterious mutations such as some transposon insertions can easily reach fixation. Moreover, as transposons quickly accumulate the probability of deletions through non-allelic homologous recombination also increases, counteracting their initial impact on genome growth (Hedges and Deininger 2007; Petrov et al. 2003). Within this context, the accordion model is an emergent property based on transposon accumulation dynamics. Importantly, the signatures of selection for an optimal genome size are not always consistent; the Chinese tree shrew has a high metabolic rate but a relatively large genome of 2.86 GB (Fan et al. 2013). This suggests that the role selection plays in driving genome size evolution is likely taxon-specific. Further, neither mechanism takes into account the underlying genome structure.

The genomic DNA of complex organisms is wrapped around nucleosomes and packaged into various conformations that regulate the access of different gene regulatory factors to their target sites. This hierarchical genome structure means that the impact and likelihood of particular mutations is highly context-specific, resulting in regional variation in both the susceptibility and tolerance to mutations. Here, susceptibility is the likelihood of a mutation occurring and tolerance is the degree to which the mutation does not adversely impact fitness. The observed accumulation patterns of DNA gain and loss events arise from the interaction of region-specific susceptibility and tolerance. For example, small (≤ 30 bp) insertion or deletion (indel) events in the human genome are correlated with recombination rate and are enriched for topoisomerase cleavage sites (Kvikstad et al. 2009, 2007). This suggests that the biological role of certain regions may cause them to be particularly susceptible to indel mutations. In the case of larger events such as transposon insertions, the prevailing model suggests that long interspersed elements (LINEs) accumulate in gene-poor regions where they are most tolerated (Gasior et al. 2007). The evolution of genome size via DNA gain and loss is not only shaped by higher order factors such as cell size and metabolic rate, but is intimately linked to the underlying genome structure.

To better characterise the molecular drivers and evolutionary impacts of DNA gain and loss, we calculated lineage-specific gain and loss rates across the human and mouse genomes. Human and mouse were chosen specifically for three reasons. Firstly, both species have well characterised genomes with highly accurate and well annotated assemblies (Lander et al. 2001; Chinwalla et al. 2002) and have both been used frequently in comparative genomic analyses resulting in many easily accessible pairwise alignment datasets available on the UCSC genome browser (Tyner et al. 2016). This makes it possible to compare them to a wide variety of outgroup species and detect genomic features that associate with DNA gain and loss. Secondly, the mouse genome is significantly smaller than the human genome, making it possible to detect a large number of lineage-specific deletion events (Chinwalla et al. 2002; Laurie et al. 2012). Finally, human and mouse genomes contain similar lineage-specific transposon families (Chinwalla et al. 2002). This means that both species share similar mechanisms for DNA gain, making it easier to compare differences between associations with other types genomic features.

For our analysis, we detected DNA gain and loss events using two distinct, yet complementary, methods from which we characterised DNA gain and loss hotspots. From this we compared the genomic distributions of our hotspots to the genomic distribution of various features associated with genome evolution and genes that participate in particular biological processes. Our results revealed that DNA gains and losses occur in different regions across autosomes, while DNA gains from both species are particularly enriched on the X chromosome where they overlap. DNA gain events generally associate with L1 accumulation and DNA loss occurs in regions associated with biological activity such as transcription and regulation. Although DNA gain and loss in human occurred mostly in different regions, they both tended to impact on the same biological processes, while in mouse DNA loss was enriched for developmental genes and DNA gain did not associate with any particular biological process.

## Materials and methods

### Net data structure and feature extraction

For feature extraction, nets were obtained from the UCSC genome browser (Kent et al. 2002, 2003). Nets are a common format for representing pairwise genome alignments. Each net contains chained blocks of aligning sequence shared between a reference and a query genome. In order for alignment blocks to be chained together their ordering must be consistent between both genomes. Often gaps between chained blocks can contain smaller chains. It is this hierarchical structuring of the highest scoring chains at the top level with lower scoring chains filling in alignment gaps that makes nets. Importantly, in the reference genome nets provide only a single layer of coverage. However, two separate nets may occasionally overlap in the query; this is usually caused by segmental duplication in the reference. These conflicts were resolved by discarding all reference nets that did not overlap nets generated from a query reference alignment. Following this filtering process, only reciprocal best hit (RBH) nets remained. In our analysis we referred to alignment blocks within a chain as ‘chain-blocks’ and the spaces between chain-blocks also within a chain as ‘chain-gaps’. The start and end coordinates in both the reference and query genome were recorded for each chain-block and chain-gap. The programs get gaps net.go and get fills net.go were used to extract all chain-gaps. Regions of chain-gaps that were overlapped by chain-blocks in lower ranked chains were discarded. Additionally, regions that were discarded as non-RBHs or fell outside of nets were plotted against synteny blocks to determine the loci hidden from our analysis in both species. Synteny data was obtained from the synteny portal (http://bioinfo.konkuk.ac.kr/synteny_portal/) (Lee et al. 2016).

### Identifying ancestral elements

Chain-blocks were extracted from all genomes identified as outgroups to human and mouse. They were combined into a single file and merged using the bedtools genomecov function with the ‘-bg’ option. This process returned a set of potential ‘ancestral elements’ along with their corresponding coverage depth. To identify false-positives and estimate the type 1 error rate, we used the genomic positions of a set of known lineage-specific repeat families in human and mouse, since lineage-specific repeat insertions should not overlap ancestral elements. The percentage overlap of our lineage-specific repeats set with ancestral elements was measured at each minimum coverage level. A similar approach was used to estimate the type 2 error rate; the type 2 error rate was estimated as the percentage of chain-blocks that did not overlap ancestral elements. To minimise our type 1 errors we selected a minimum coverage depth threshold independently for both hg19 and mm10, where nucleotide positions with coverage depth below the threshold were not considered as ancestral elements. The basis for this approach was that nucleotide positions in our reference genomes that aligned to a large number of outgroup species were highly likely to share ancestry with those species. In contrast, nucleotide positions in our reference genomes that aligned to very few outgroup species were likely errors caused by spurious alignments between complex regions that are difficult to map. Importantly, reductions in our type 1 error caused an increase in our type 2 error. Therefore, we chose the highest possible minimum coverage threshold, where the gain in the cumulative proportion of type 1 errors from lower threshold values was greater than the gain in proportional increase of type 2 errors.

### Identifying recent transposon insertions

For both hg19 and mm10, genomic coordinates for transposons were obtained from the Repeat Masker database (Smit et al. 2015). Based on their overlap with chain-blocks or ancestral elements, individual transposons were classified as either recent or ancestral. In addition to this, the percent divergence from consensus family sequence and the proportion of total sequences of transposon family members that overlapped ancestral elements or chain-blocks were calculated. This data was then used in linear discriminant analysis to build a transposon family classifier. Our classifier was trained using the original individual transposon classifications. After training, entire families were classified as either recent or ancient using the family-wise means of the feature values. Finally, transposons from families classified as recent but overlapping gaps between reference and query were classed as lineage-specific insertions.

### Gap annotation and placement

Chain-gaps extracted from nets were annotated as either DNA gain or DNA loss based on two distinct yet complementary annotation methods; the recent transposon-based method and the ancestral elements based method. The ancestral element-based method infers the ancestral state of a gap. For example, an mm10 gap overlapping an ancestral element would be annotated as an mm10 loss, whereas the same gap not overlapping an ancestral element would be annotated as an hg19 gain. The recent transposon-based method instead identifies DNA gains. In this case an mm10 gap overlapping a recent transposon would be annotated as an hg19 gain, while an mm10 gap not overlapping a recent transposon would be annotated as an mm10 loss.

After all chain-gaps between a reference and query were annotated in both genomes, the remaining non-aligning sequences were ‘placed’ in the genomes they were absent from. This process is referred to as ‘gap placement’ and is performed on the non-aligning sequence of chain-gaps that remain in the reference genome after a reference query alignment. These non-aligning reference sequences are absent from the query and are either the result of DNA gain in the reference or DNA loss in the query. Using the coordinate mappings of the 5’ and 3’ adjacent chain-blocks of each chain-gap, the non-aligning reference sequence of a chain-gap is inserted into the query genome at the corresponding position, where placed gaps are oriented relative to the genome they are placed in. Importantly, gap placement begins by placing chain-gaps at the bottom chain level of nets and ends by placing chain-gaps at the top chain level. This process ensures that non-aligning sequence in overlapping chain-gap annotations caused by hierarchical structure of nets are only placed once. Once the corresponding position of a gap has been identified, the downstream query coordinates are incremented by the size of the annotated chain-gap being placed. This creates a synthetic genome consisting of DNA gains and losses that occurred across both the reference and query lineages. The total length of our synthetic genomes is equal to the total length of the query genome and the total length of annotated chain-gaps from the reference. Finally, the synthetic genomes were segmented at a window size of 200kb into distinct genomic bins where the total size of each gap annotation was tallied. Genomic bins with less than 150 kb that did not belong to assembly gaps or non-RBH regions were discarded. Importantly, our decision to use a synthetic genome meant that placed chain-gaps larger than our window size would spread across window boundaries, ensuring that genomic bins would contain no more than 200 kb of sequence.

### Hotspot identification

Hotspots for reference gain, reference loss, query gain and query loss in both hg19 and mm10 were identified using the Getis-Ord local statistic found in the R package ‘spdep’ (Bivand et al. 2013; Bivand and Piras 2015). The Getis-Ord local statistic for genomic bin *i* is calculated as:

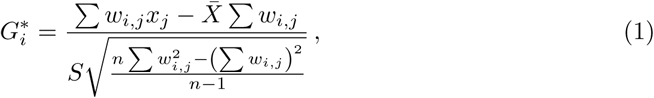

where *x*_*j*_ is the number of bp belonging to a particular gap annotation within bin *j*, *w*_*i,j*_ is the atial weight between bin *i* and *j*, *n* is the number of bins for a particular genome, 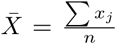 and 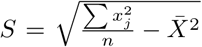 (Getis and Ord 1996). For the neighbourhood weight matrix *W*, *w*_*i,j*_ was given a spatial weight of 1 if bin *i* and bin *j* were considered neighbours. For bin *i* and *j* to be considered neighbours bin *j* had to be within 600 kb of bin *i*. After calculating 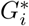 for each bin and each gap annotation in both genomes, all 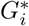 values were converted to P-values and adjusted for multiple testing using the false discovery rate (FDR). Bins were only considered hotspots if their 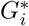 was > 0 and had a FDR < 0.05.

### Obtaining genomic features

A set of genomic features was obtained from a range of sources to identify factors potentially driving DNA gain and loss. GC content was calculated as the proportion of chain-blocks per bin using the hg19 and mm10 Biostrings-based genome R packages (Team TBD 2014a,b; Pages 2017). CpG islands for both hg19 and mm10 were obtained from the UCSC genome browser (Tyner et al. 2016). DNaseI hypersensitivity (DNaseI HS) peaks for hg19 were obtained from UCSC as part of the DNaseI master track (http://hgdownload.cse.ucsc.edu/goldenpath/hg19/encodeDCC/wgEncodeAwgDnaseMasterSites/). The master track was generated by combining DNaseI HS sites from across 125 cell lines produced by the University of Washington and Duke University ENCODE groups (ENCODE Project Consortium et al. 2012). The Individual cell line data can be located using the accessions GSE29692 and GSE32970. DNaseI HS peaks for mm10 were obtained from UCSC as individual samples mapped to mm9 (https://genome.ucsc.edu/cgi-bin/hgFileUi?db=mm9\&g=wgEncodeUwDgf). Individual peaks from each sample were merged into a single file, creating a single set of DNase1 HS peaks. The merged mm9 peaks were then converted to the mm10 assembly using the UCSC liftover tool (Hinrichs et al. 2006). Mouse DNaseI HS peaks were generated using DNaseI digital genomic foot-printing performed by the University of Washington ENCODE group (ENCODE Project Consortium et al. 2012). This data can also be obtained using the accession GSE40869. Recombination rates for human were identified as part of the HapMap project (ftp://ftp.ncbi.nlm.nih.gov/hapmap/recombination/2011-01_phaseII_B37/)(International HapMap Consortium et al. 2007). However, recombination hotspots were only available for earlier phases of the HapMap project (ftp://ftp.ncbi.nlm.nih.gov/hapmap/recombination/2006-10_rel21_phaseI+II/hotspots/). The hotspots were initially mapped to hg17 and then converted to hg19 coordinates using the UCSC liftover tool. Recombination hotspots were identified using the methods outlined in Winckler et al. (2005) and McVean et al. (2004). Recombination rates and hotspots in mouse were calculated in mm9 based on two separate datasets (Brunschwig et al. 2012; Kirby et al. 2010; Yang et al. 2011). They were converted to mm10 using the UCSC liftover tool. Importantly, recombination data was only available for mouse autosomes. During enrichment tests this was taken into account by removing the sex chromosomes from the sample space. Exons and introns for both hg19 and mm10 were extracted from UCSC genome annotations available from TXDB R packages (Carlson 2015, 2016; Lawrence et al. 2013). Retrotransposon coordinates for hg19 and mm10 were obtained from the Repeat Masker database (http://www.repeatmasker.org/genomicDatasets/RMGenomicDatasets.html) (Smit et al. 2015). The Repeat Masker version used for hg19 and mm10 was open-4.0.5 with repeat library 20140131. Retrotransposons were sorted into the following categories: ancient elements, ancestral L1s, lineage-specific L1s and lineage-specific SINEs using prefixes for families of known lineage-specific and ancestral activity (Giordano et al. 2007). Ancient elements were identified by the class names ‘SINE/MIR’ and ‘LINE/L2’. Ancestral L1s were identified using the family name prefixes ‘L1ME’, ‘L1MD’, ‘L1MC’, ‘L1MB’ and ‘L1MA’. Human lineage-specific L1s were identified using the family name prefixes ‘L1PB’, ‘L1PA’ and ‘L1HS’. Mouse lineage-specific L1s were identified using the family name prefixes ‘Lx’, ‘L1Md’, ‘L1 Mus’, ‘L1 Mur’ and ‘L1 Mm’. Human lineage-specific SINEs were identified using the family name prefix ‘Alu’. Mouse lineage-specific SINEs were identified using the family name prefixes ‘PB’, ‘B1’, ‘B2’, ‘B3’ and ‘B4’. Lamina associated domains (LADs) for hg19 were obtained from the UCSC genome browser (http://hgdownload.soe.ucsc.edu/ goldenPath/hg19/database/laminB1Lads.txt.gz) (Guelen et al. 2008). LADs for mouse were constitutive across several samples and were obtained using the accession GSE17051, they were converted from mm9 assembly to mm10 assembly using the UCSC liftover tool (Peric-Hupkes et al. 2010). For each feature, except recombination rate, the per 200 kb coverage level for each bin was calculated. For recombination rate the mean rate per bin was used.

### Genomic feature enrichment

Feature enrichment was detected on the basis of a permutation test. For each feature and hotspot in both hg19 and mm10, a background distribution was generated by calculating the difference in means between a set of resampled hotspot and non-hotspot bins 10,000 times, resampling was performed without replacement. The background distribution was then used to convert the differences in means between observed hotspot and non-hotspot bins into a Z-score to allow standardisation between features and gap annotations and provide the direction of the association.

### GO term enrichment analysis

Gene ontology (GO) term enrichment was calculated using the topGO package in R (Alexa and Rahnenfuhrer 2016). Genes within each hotspot region were independently tested against the genomic background. For enrichment, the Fisher test was used in combination with four separate algorithms: the classic algorithm treats each term independently whereas the elim, weight and parent-child algorithms factor in the GO inheritance structure (Alexa et al. 2006; Grossmann et al. 2007; Ashburner et al. 2000); the elim algorithm removes all genes annotated to a significantly enriched GO term from all of the terms ancestors; the weight algorithm behaves similarly, instead of removing genes from the ancestors of enriched GO terms, it creates a more subtle effect by reducing the weight of genes annotated to the ancestors of enriched GO terms (Alexa et al. 2006); for the parent-child algorithm, the enrichment score for a particular term takes into account the probability a random set of genes of the same size contains the same exact parents (Grossmann et al. 2007). Because these algorithms adjust the enrichment probabilities they obviate the need to account for multiple testing (Alexa and Rahnenfuhrer 2016).

### Software and data analysis

All statistical analyses were performed using R including the packages GenomicRanges, RMySQL, dplyr and Bioconductor (R Core Team 2015; Lawrence et al. 2013; Ooms et al. 2016; Wickham and Francois 2015; Gentleman et al. 2004). Code used to perform analyses can be found at: https://github.com/AdelaideBioinfo/regionalGenomeTurnover.

## Results

### Detecting DNA gain and loss events

Across genomes and throughout evolution DNA is frequently gained and lost by the processes of insertion and deletion. To identify individual events and quantify DNA gain and loss at a regional level in hg19 and mm10, we obtained pairwise alignment data between both genomes in the form of nets from the UCSC genome browser (methods) (Tyner et al. 2016; Kent et al. 2003). By taking advantage of the data’s hierarchical structure we were able to estimate DNA gain and loss in regions that have undergone rearrangements. We processed our data in three distinct steps; 1) extract features (Fig. 1a), 2) annotate gaps (Fig. 1b-c) and 3) place gaps (Fig. 1d).

**Figure 1.**
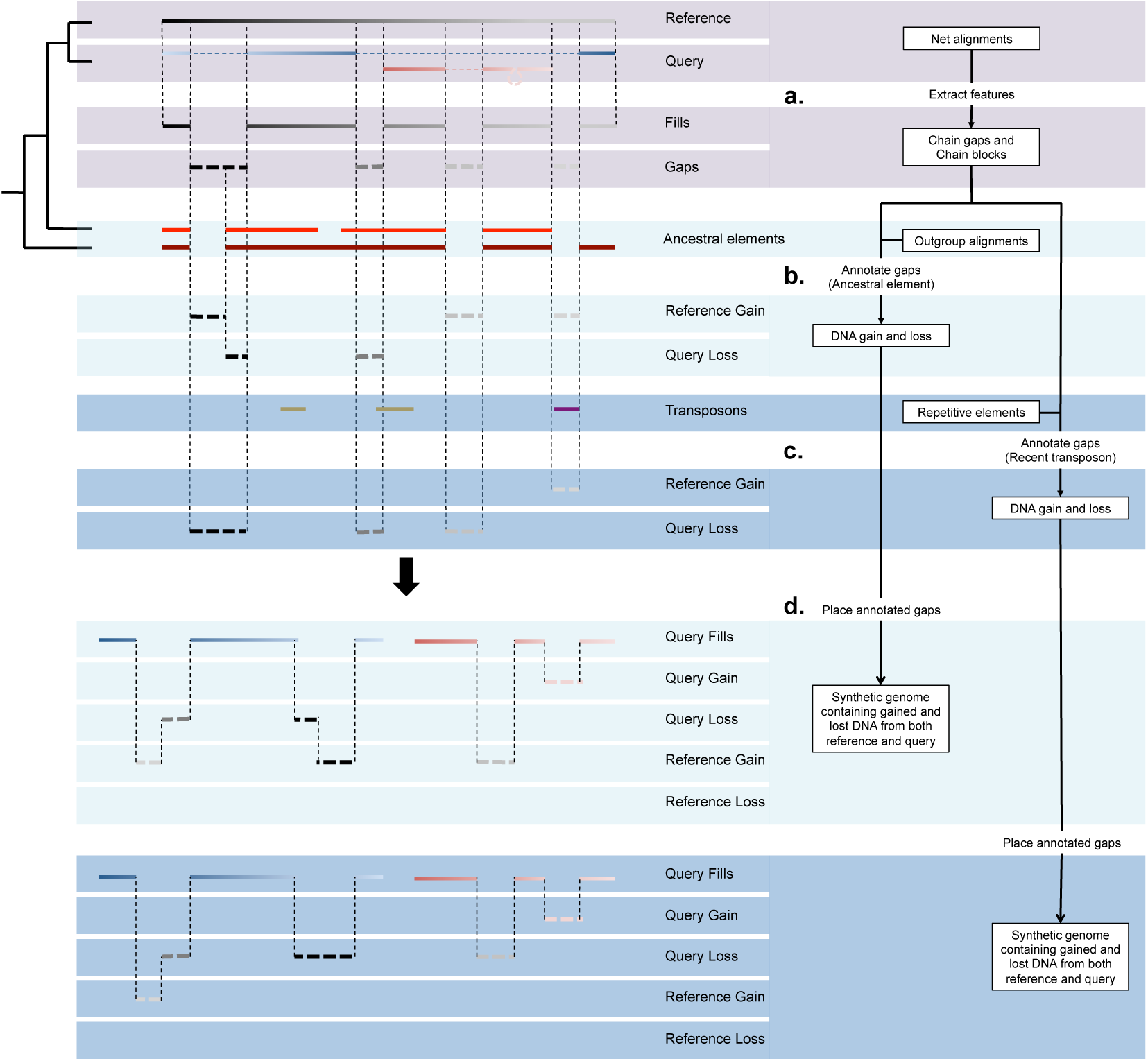
Detecting DNA gain and loss events between two species. Chain-gaps and chain-blocks are extracted from nets between reference and query (a). The resulting chain-gaps are essentially sequences from the reference genome that do not align to anything in the query genome. Chain-blocks are extracted from nets between reference and outgroup species as ancestral elements. Ancestral elements are then used to annotate chain-gaps as either gain or loss (b). Chain-gaps are annotated as query loss if they overlap ancestral elements or as reference gain if they do not. This is the ancestral element method for annotating gaps. The recent transposon method instead uses transposons classified as recent or ancestral to annotate gaps (c). Transposons are extracted from Repeat Masker files containing various classes of repetitive elements. Chain-gaps are annotated as reference gain if they overlap recent transposons or as query loss if they do not. After gaps are annotated they are placed within each genomic background creating a synthetic genome (d). Annotated chain-gaps are placed according to the edge coordinates of their adjacent chain-blocks within the same chain. Shown in the final two panels are chain-gaps extracted from the reference placed within the query genome. The different colours of the query chain-blocks show that gap annotations in the reference are placed on different chromosomes in the query. Differences in annotations are the results of conflicting information either resulting from incorrect identification of ancestral elements or recent transposons.

For step 1, chain-gaps and chain-blocks were extracted from nets considering only chain-gaps of at least 10 bp in size (Fig. 1a) (methods). Our approach allowed us to keep track of each feature’s position in both the reference and query genome. This is especially important since it is not possible to identify deletions when the corresponding coordinates between species are lost. After extracting features we found that approximately 111 Mb of hg19 and 174 Mb of mm10 were not contained within nets (Table 1). Alignment gaps that didn’t belong to any nets in human and mouse tended to overlap regions between two conserved synteny blocks (Fig. S1-S2). With the remaining features extracted from hg19 and mm10, we used the corresponding coordinates between reference and query to identify features that were reciprocal best hits (RBHs). This removed features in the reference genome that mapped to similar locations in the query, which are likely the result of segmental duplication. After filtering out non-net and non-RBH regions, 1014.3 Mb of chain-blocks and 1465.8 Mb of chain-gaps remained in hg19, and 994.4 Mb of chain-blocks and 1191.5 Mb of chain-gaps remained in mm10 (Table 1). Since our processed nets for each genome are supposed to only contain RBH features, it is expected that the coverage of chain-blocks should be equal between hg19 and mm10. To determine the source of this discrepancy, we analysed the number of chain-gaps bellow our minimum size cut off and found that when these were taken into consideration the difference in chain-block size was reduced to approximately 1 Mb.

**Table 1.** Processing of net files. Sizes of genomic regions are measured in Mb unless otherwise specified.

| Genomic regions | hg19 | mm10 |
| --- | --- | --- |
| Sequenced genome | 2897.0 | 2653.0 |
| Gaps outside of nets | 111.1 | 174.0 |
| Non-RBH chains | 306.1 | 293 |
| Ancestral elements | 1726.0 | 1021.0 |
| Remaining chain-blocks | 1014.3 | 994.4 |
| Remaining chain-blocks ∩<br>ancestral elements (%) | 94.2 | 85.2 |
| Remaining chain-gaps | 1465.8 | 1191.5 |

Next, for step 2 we annotated chain-gaps as either lineage-specific DNA gain or DNA loss. To annotate gaps we used two complementary methods, an ancestral elements-based method and a recent transposon-based method. The ancestral element-based method uses outgroup species to annotate gaps by inferring their ancestral state (Fig. 1b). For example, if a particular sequence between a reference and outgroup is conserved but presents as a gap in the query it is likely that this sequence was lost from the query. Alternatively, if this particular sequence in the reference presents as a gap in both the query and the outgroup it is likely that this sequence was instead gained in the reference. An important consideration for identifying ancestral elements is the type 1 (false positive) and type 2 (false negative) error rates, where type 1 errors are lineage-specific regions annotated as ancestral elements and type 2 errors are ancestral regions annotated as lineage-specific. To reduce our type 2 error rate we obtained the genomes of a large range of human and mouse outgroup species from the UCSC genome browser (Table S2). Across all of our outgroup species we extracted all the chain-blocks and merged overlapping intervals to create our ancestral elements. This strategy increased the chance of finding ancestral DNA in our reference that may have been lost in one or more of our outgroup species. For both hg19 and mm10 we found that total genome coverage of ancestral elements reached asymptotic levels at approximately 18 outgroup species (Fig. S3). However, this strategy also came with the trade-off of increasing our type 1 error rate. To control error rates we measured how type 1 and type 2 errors responded to changes in coverage depth of outgroup chain-blocks at each position in hg19 and mm10 (Fig. S4). Based on these results we annotated human ancestral elements at an outgroup coverage depth ≥ 6 and mouse ancestral elements at an outgroup coverage depth ≥ 4 (Fig. S4). This strategy removed > 85% ancestral elements overlapping known lineage-specific repeats in mouse and > 95% of ancestral elements overlapping known lineage-specific repeats in human. For remaining chain-blocks, we found that 94.2% in human and 85.2% in mouse were supported by our annotated ancestral elements (Table 1). Our very low error rate in human indicates that we were able to accurately determine the amount of mm10 DNA loss and hg19 DNA gain. However, our error rates in mm10 suggest that ancestral regions alone are insufficient to accurately estimate hg19 DNA loss and mm10 DNA gain.

To complement and overcome potential shortcomings of the ancestral element-based method of estimating DNA gain and loss, we adopted a recent transposon-based method. We identified transposon families with lineage-specific activity and used them to annotate gaps as lineage-specific DNA gain or loss (Fig. 1c). For example, recent transposon sequences in hg19 that overlap gaps in mm10 are annotated as hg19 gains, where ancestral transposon sequences in hg19 that overlap gaps in mm10 are annotated as mm10 losses. This approach has been used previously to identify DNA loss in the mouse and human lineages (Chinwalla et al. 2002; Hardison et al. 2003).

In order to annotate gaps using the recent transposon method, we first had to identify transposon insertions that occurred after mouse and human diverged from their common ancestor. Because transposon families have undergone distinct bursts of activity at particular points in time, we decided to classify transposon families as either ‘recent transposons’ or ‘ancestral transposons’, and use members of those respective classifications to annotate our chain-gaps. The main challenge in this approach is identifying lineage-specific activity of transposons. Generally, transposon families are considered to be ancestral transposon families when they are shared between two species. However, there is a possibility some ancestral transposon families may have been active during the period of human and mouse divergence and continued replicating in each lineage independently. This means families that would have been otherwise classified as ancestral transposons may have actually undergone varying amounts of lineage-specific transposition.

To overcome the problem of misclassifying the activity of otherwise ancestral transposon families, we used linear discriminant analysis to build a transposon family classifier for both human and mouse. We initially obtained transposon coordinates from the Repeat Masker database and classified individual transposons as ‘ancestral transposons’ if they overlapped ancestral elements or chain-blocks and as ‘recent transposons’ if they did not. Next, we trained our classifier using two separate variables. The first variable was each transposon’s percent divergence from their family consensus sequence, often used as an indicator of transposon age (Kapitonov and Jurkal 1996; Smit et al. 1995). The second variable was the proportional overlap between each transposon family and ancestral elements or chain-blocks as measured by bp coverage. After training we used our classifier to group each family based on the family-wise means for the variables above (Fig. S5). We identified 656 recent human transposon families and 689 recent mouse transposon families. Our results suggest that at least 176 families were active during human and mouse divergence leading to a mixture of both ancestral and lineage-specific insertions (Table S1). Moreover, the percent divergence of these families is consistent with transposon activity occurring after the evolution of ancestral transposons and prior to the evolution of lineage-specific transposons (Fig. S6). Surprisingly, we also identified some transposon families that were not shared between human and mouse, and yet were annotated as ancestral. However, these families were usually small and together they covered less than 1 Mb of their respective genomes (Table S1). In addition, our results for mm10 indicate potential drawbacks in using the ancestral element-based method for annotating gaps; percent divergence from consensus for some recent transposon families is similar to ancestral transposon families. While this is consistent with an elevated rate of substitution in the rodent lineage, it suggests that a large number of regions in mm10 that share ancestry with our outgroup species may have diverged beyond the alignment threshold (Fig. S5). Collectively, these results demonstrate the difficulty of identifying recent transposon insertions based on family name alone. For this reason we decided to annotate chain-gaps using our newly classified recent transposon families, which were classified using a combination of family-wide and transposon-specific factors in conjunction with comparative genomic approaches.

Using both the ancestral element and recent transposon based methods, we annotated a large number of chain-gaps with varying levels of consistency. In hg19, both methods were largely consistent in identifying human-specific DNA gains and mouse-specific DNA loss. However, in mm10 there was less agreement between the methods; while the majority of mouse lineage-specific DNA gains identified by both methods tended to overlap, the majority of human lineage-specific DNA loss did not (Table 2). This is mostly likely due to limitations for detecting ancestral elements in mm10. We found that only 85% of mm10 chain-blocks were supported by ancestral elements as opposed to 95% in hg19 (Table 1), suggesting that many ancestral elements were not identified using our outgroup species. This is a key weakness in our approach; if there is an underlying error for detecting human DNA loss in mm10, it means that we would also be overestimating the amount DNA gain in mm10. However, by using two distinct yet complementary methods, we are able to identify potential sources of error and estimate their impact. One explanation for missing ancestral elements may be that DNA gain and loss events that occurred in either the mouse or human clade overlap DNA gain and loss events that occurred across a large number of our outgroup species. However, as stated above, nucleotide divergence rates may also play a role. Some regions in mm10 may have diverged so much that it is impossible to perform a pairwise alignment with our outgroup species. Despite the above mentioned inconsistencies between the methods in mm10, it is clear that the amount of DNA loss in human is much smaller than the amount of DNA loss in mouse and the amount of DNA gain for both. The difference in loss rates for human and mouse is mostly consistent with a high deletion rate in the mouse genome that has caused it to shrink in size since divergence with human (Chinwalla et al. 2002; Laurie et al. 2012).

**Table 2.**
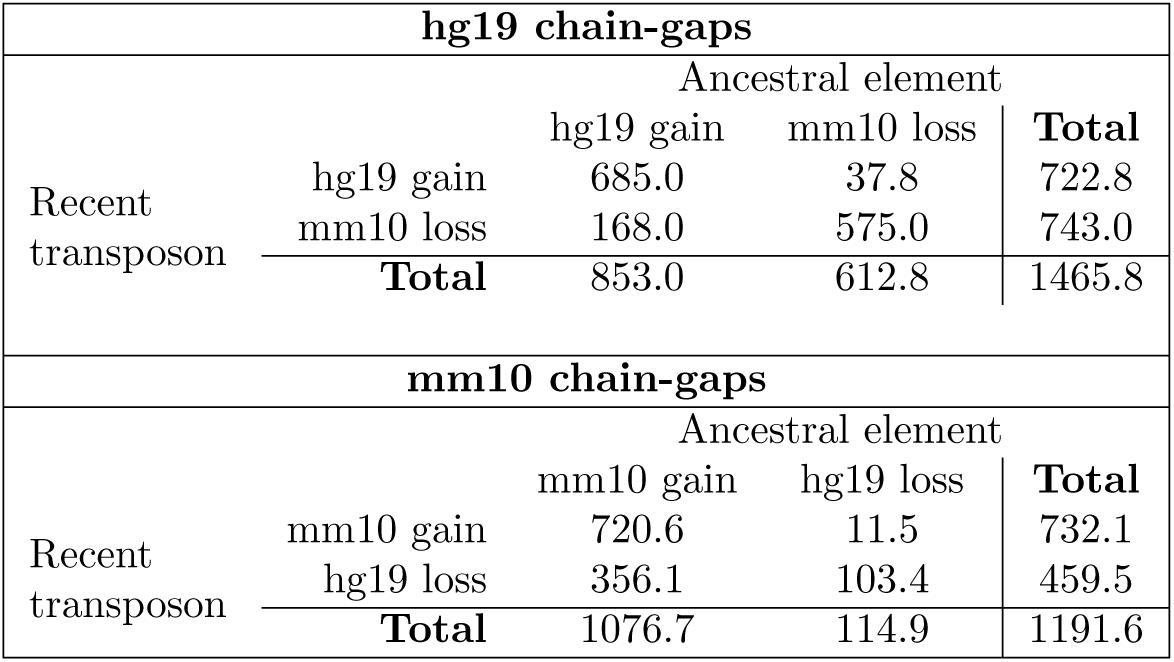
hg19 and mm10 gap annotation. Chain-gaps were annotated using both the ancestral element and recent transposon method. Each number represents gap annotations in Mb.

To further characterise the results from each method we compared the length distributions of their gap annotations. For DNA gain events in hg19 and mm10, the ancestral element method displayed a much higher frequency of small elements than the recent transposon method. This may be caused by spurious alignments between similarly structured recent transposons found in reference and outgroup species, effectively separating the annotation gain events into smaller pieces. Moreover, the recent transposon method identified much higher frequencies of DNA gain events that correspond to full length consensus sequences of known transposon families (Fig. 2a-2b). Conversely, the length distributions for DNA loss events identified by each method were much more similar, especially in mm10. In hg19 the frequency of events detected by the ancestral element method were much lower than those detected by the recent transposon method (Fig. 2c-2d). This is consistent with the low number of ancestral elements in the mouse genome. However, the high level of consistency for both methods in identifying hg19 DNA gain and mm10 DNA loss where there is good support for outgroup species is highly encouraging. It indicates that the recent transposon method is a reasonably effective method in identifying DNA gain and loss in species where it is difficult to detect ancestral elements. Consistent between both methods is size distribution difference between DNA gain and loss. DNA gain events are mostly over 100 bp in length while DNA loss events are mostly under 100 bp.

**Figure 2.**
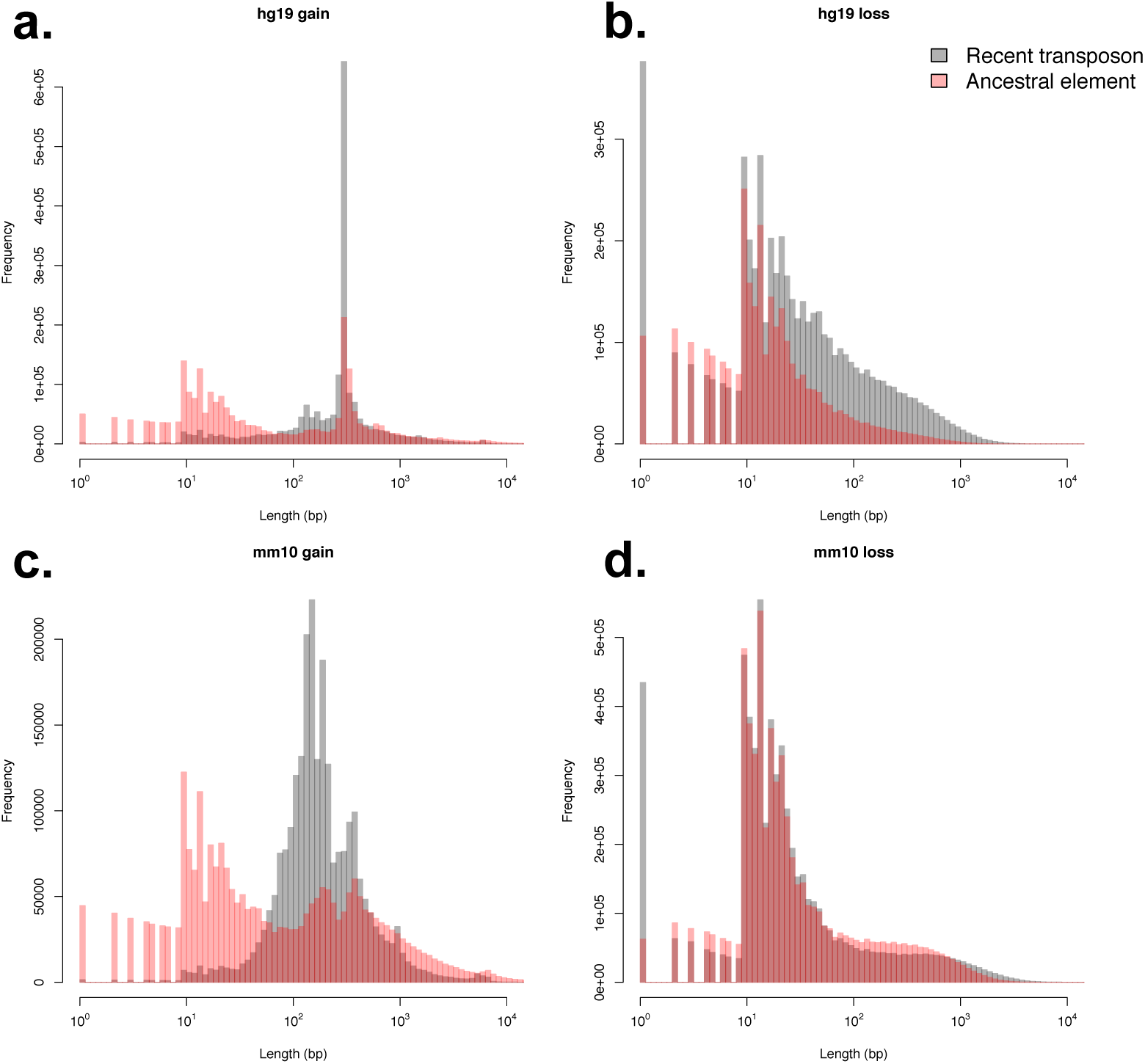
Length distributions of identified DNA gain and loss events. hg19 gain (a), mm10 gain (b), hg19 loss (c) and mm10 loss (d) events were identified using both the recent transposon and ancestral element method. Peaks for hg19 and mm10 gain, especially those detected by the recent transposon method, correspond to know lengths of transposon families.

In both hg19 and mm10 we annotated a large number of gain and loss events using two distinct methods. However, to measure the total amount of DNA turnover at particular loci, gaps annotated in a query genome needed to be mapped to a reference genome. Hence, gap annotations were placed using the reference and query coordinates we extracted from our nets in step 1 (methods) (Fig. 1d). To account for the placement of gaps from one genome into another, we adjusted the genomic coordinates at the target loci, resulting in a synthetic genome for both species (methods). Each synthetic genome contains both hg19 and mm10 annotated gaps in either an hg19 or mm10 genomic background. Finally, our resulting dataset consists of 4 synthetic genomes; mm10 with gap annotations based on the ancestral element method, mm10 with gap annotations based on the recent transposon method, hg19 with gap annotations based on the ancestral element method and hg19 with gap annotations based on the recent transposon method. Collectively, these results demonstrate that it is possible to identify locations for the majority of DNA gain and loss events since human and mouse divergence. Using our identified DNA gain and loss events it is possible to characterise genome-wide patterns of DNA gain and loss and to begin to determine how DNA turnover may impact on mammalian genome evolution.

### Genome-wide characteristics of DNA gain and loss

Genome size evolution in mammals follows an accordion model, where DNA gain is counteracted by DNA loss to maintain a relatively constant genome size (Kapusta et al. 2017). To characterise how DNA gain and loss interacts with genome structure, we used our synthetic genomes to analyse the genomic distribution of DNA gain and loss events in hg19 and mm10. We began by segmenting synthetic genomes into 200 kb non-overlapping bins and tallying the total bp coverage of each type of gap annotation. Bins with less than 150 kb of DNA not belonging to RBH nets were removed and our tallies were normalised to reflect DNA gain and loss amounts per 200 kb. Because gap annotations from both species can be placed within a single genome, we are able to directly compare their genomic distributions.

Using our binned synthetic genomes we compared the variation and average amount of regional DNA gain and loss identified using each method. Our results showed that variation in regional DNA gain or loss was reasonably consistent across both methods (Fig. 3). For DNA gain this was also quite large, in 200 kb genomic bins the amount of DNA gain in human and mouse spanned a range greater than 70 kb, indicating that some regions underwent much greater levels of DNA gain than others. While bin-wise variation in gain and loss rates was consistent across methods, the average amount of DNA turnover was not. This makes it difficult to reliably calculate the regional amount of DNA turnover or genome growth. However, despite these inconsistencies, bin-wise levels of DNA gain and loss were highly correlated across all cases, with the exception of hg19 DNA loss (Fig. 3a, S7-S8). Following this, we investigated regional DNA gain and loss dynamics by identifying DNA gain and loss genomic hotspots. Hotspots were identified by calculating 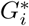 for each bin (methods). We converted our 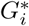 values to P-values and calculated the false discovery rate (FDR). Bins whose 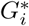 was positive with FDR < 0.05 were considered hotspots. Hotspots were identified for each type of gap annotation found using both gap annotation methods in both synthetic genomes. We found that the size of the hotspot overlap between each gap annotation method for hg19 gain, mm10 gain and mm10 loss was larger than the sum of non-overlapping hotspots (Fig. 3b). Using the hotspot intersect between gap annotation methods, we further characterised regional variation of DNA gain and loss across hg19 and mm10. For the remainder of the analysis the terms ‘DNA-gain hotspots’ and ‘DNA-loss hotspots’ refer to the hotspot intersect between each gap annotation method, except for hg19 DNA-loss hotspots which instead refer to hg19 DNA-loss hotspots identified through the recent transposon method. For mm10 DNA loss, mm10 DNA gain and hg19 DNA gain, the intersect was used as it provided a sample of genomic regions where regional DNA gain and loss dynamics were highly supported by both methods. For hg19 DNA loss we used hotspots that were identified using the recent transposon method because the ancestral based method was shown to largely underestimate the total amount of ancestral DNA.

**Figure 3.**
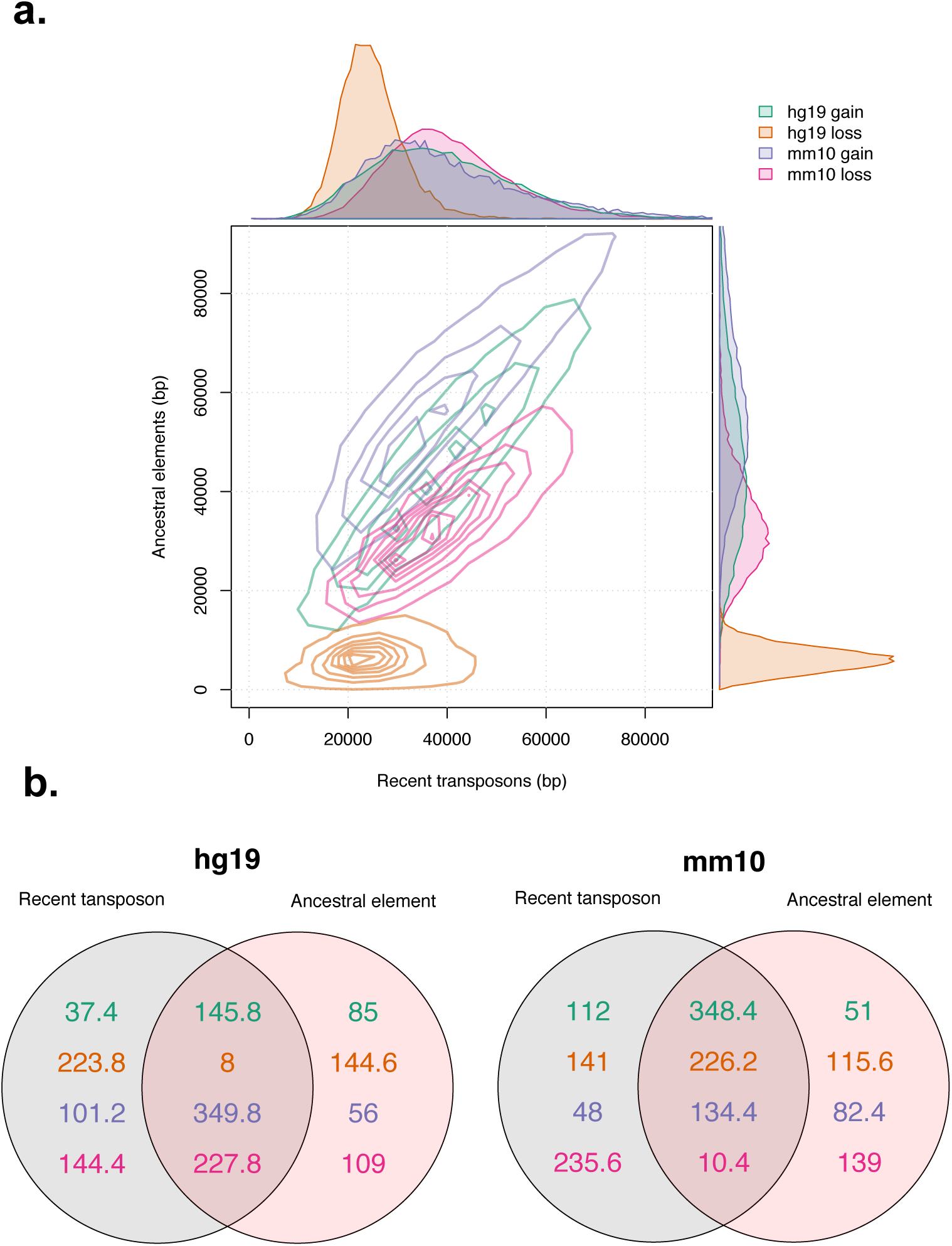
Comparison of gap annotation methods in binned synthetic genomes. Amount of DNA gain and loss per 200 kb in each bin for both hg19 and mm10 (**a**). For each gap annotation, contour lines begin at a 2D kernel density estimate of 2^−10^ and increase at regular intervals of 4^−10^, except for hg19 which increase at regular intervals of 1.6^−9^. Sizes of regions in Mb identified as hotspots for DNA gain or loss using the 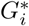 statistic in each genome (**b**).

### Regional patterns of DNA gain and loss indicate lineage-specific divergence

The accordion model of genome evolution suggests DNA gain and loss is largely balanced across the entire genome. Whether the individual events are balanced at the local scale remains unknown. We analysed the genomic distribution of hg19 and mm10 gain and loss hotspots by focussing on the within species overlap and the across species overlap. The within species overlap was designed to investigate whether DNA gain and loss is balanced on a regional level, indicating that despite large amounts of DNA turnover, local genome structures stay intact. The across species overlap was designed to investigate whether DNA gain and loss associated with lineage specific divergence in genome architecture. We found that almost 4% of human loss hotspots overlapped human gain hotspots and approximately 6% human gain hotspots overlapped human loss hotspots (Fig. 4,S9). These results showed that DNA gains and losses in human at a regional scale have occurred independently. Conversely, less than 1% of gain and loss hotspots in mouse overlapped each other, with a significant negative association. These results suggest that regional DNA gain and loss in both species is largely unbalanced. For the across species comparison, we found significant levels of overlap between DNA-loss hotspots and negative associations between all other hotspot types at varying levels of statistical significance depending on genomic background. This demonstrates that DNA loss dynamics in both hg19 and mm10 share some degree of conservation while DNA gain dynamics are mostly lineage-specific, suggesting that the acquisition of new DNA may be driving lineage-specific divergence of genome structure.

**Figure 4.**
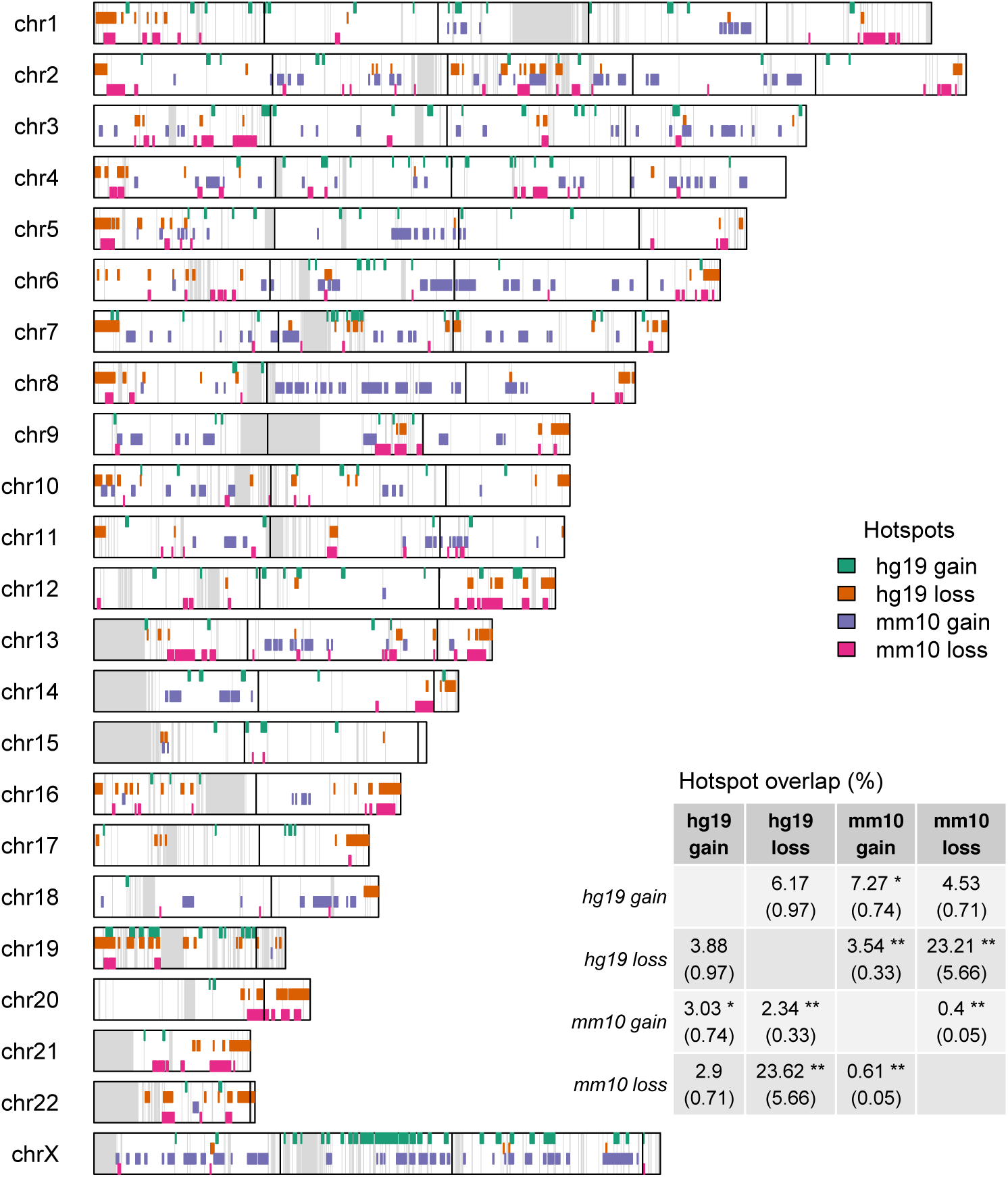
Genomic distribution of gain and loss hotspots for hg19 and mm10 plotted against hg19 synthetic genome. Grey regions indicate bins with <150 kb of RBH nets and black vertical lines represent 50 Mb on non-synthetic genome. Inset table represents percent overlap of gain and loss hotspots. The percentages were calculated using the hotspots labelled in each row as the denominator. ‘*’ and ‘**’ represent p-values below 0.05 and 0.01 respectively based on the Fisher statistic. The odds ratio for each fisher test is reported within the brackets. An odds ratio above 1 represents a positive association and an odds ratio below one represents a negative association.

To further characterise the distribution of hg19 and mm10 gain and loss hotspots, we plotted them against both genomic backgrounds. hg19 and mm10 gain hotspots were most enriched on chromosome X (Fig. 4,S9). This is consistent with chromosome X as a hotspot for L1 insertion, a particularly large transposon with high levels of lineage specific activity that contributes to X inactivation (Chow et al. 2010). For gain and loss hotspots themselves, hg19 gain hotspot regions were much more dispersed than other types of hotspot region (Fig. 4,S9). Since DNA loss across both species overlaps significantly, this adds to the lineage-specific behaviour of DNA gain dynamics, where regional DNA gain in mouse is more concentrated than in human. Interestingly, DNA loss hotspots in the hg19 genomic background appear more concentrated towards telomeres, suggesting that chromosomal location may play a role in DNA loss dynamics (Fig. 4). However, it is worth noting that this observation did not occur in the mm10 genomic background (Fig. S9). One explanation is that telomeres in mouse are quite recent as mouse chromosomes have undergone a high frequency of breakage and fusion events since divergence from a common ancestor (Murphy et al. 2005). Together, our results demonstrate that regional lineage-specific DNA gain and loss dynamics are relatively context-specific.

Next, we examined whether gain and loss hotspots were correlated with a range of genomic features. The genomic features we analysed are non-randomly distributed and known to play various roles in genome biology. By investigating their association, we may begin to develop insight into the molecular drivers of DNA turnover. To measure the correlation between genomic features and particular gap annotations we performed feature enrichment analysis with 10,000 permutations (methods). The analysis was performed for both mm10 gain and loss and hg19 gain and loss in both the genomic backgrounds. Using both genomic backgrounds we were able to analyse the genomic features from regions in a query genome that have been deleted from a reference. We specifically chose genomic features that could be found in both genomes as indicators for distinct aspects of genome biology. Intron density, exon density, DNaseI hypersensitivity (DNaseI HS) peaks, CpG islands, GC content and lamina-associated domains (LADs) are all indicators of genome activity (ENCODE Project Consortium et al. 2012; Tyner et al. 2016; Guelen et al. 2008; Peric-Hupkes et al. 2010). Most of these features, excluding LADs, are associated with gene dense areas and are linked to their expression or regulation (Thurman et al. 2012). LADs themselves are instead associated with gene-poor regions and gene silencing (Guelen et al. 2008; Peric-Hupkes et al. 2010). We also investigated various groups of transposons whose genomic distributions have been previously characterised and used to investigate genome-wide DNA gain and loss rates. Lineage-specific L1s and SINEs are both major sources of DNA gain via retrotransposition, they both also have distinct accumulation profiles that are similar across both species (Chinwalla et al. 2002). Lineage-specific L1s tend to accumulate in gene-poor regions while lineage-specific SINEs accumulate in gene rich regions. Ancestral L1s, and ancient elements (MIRs and L2s) have been used previously to indicate levels of DNA loss. Since these elements inserted prior to species divergence, they both provide signatures of ancestral DNA. Differences in the numbers of these elements in similar regions across species can indicate DNA loss (Chinwalla et al. 2002; Laurie et al. 2012). Finally, we investigated the genomic distribution of recombination hotspots and genome-wide profiles of recombination rates (International HapMap Consortium et al. 2007; Brunschwig et al. 2012). We considered recombination as an indicator of genome instability, as meiotic recombination increases the potential for heritable genomic rearrangements (Berg et al. 2010). Importantly, it is worth noting that recombination hotspots and recombination rates in mm10 are autosomal only. This was due to limited data availability for mouse.

Among our features we observed distinct profiles for DNA gain and loss that were largely consistent across both genomes. For DNA loss from both genomes and in both genomic backgrounds we found a strong positive associations with indicators of gene-rich/active genomic regions. This is surprising as biologically active genomic regions are likely to contain many important functional elements. However, it has recently been shown that these regions are particularly prone to genomic instability leading to evolutionary genomic rearrangements (Berthelot et al. 2015). This also suggests the DNA loss is linked to an open chromatin state as it is strongly negatively associated with LADs. In the hg19 genomic background we also found that ancient elements were positively associated with mm10 DNA loss. While ancient elements have been used as indicators of DNA loss, we did not expected they would be quite so strongly associated with it. Moreover, in hg19 ancient elements are negatively associated with DNA loss and have been predicted to play important roles in gene regulation (Kamal et al. 2006). In addition, the high DNA loss rate in these regions may lead to overestimates of the genome-wide DNA loss rate in mouse, as these elements have previously been used as markers for calculating deletion rates (Lander et al. 2001; Chinwalla et al. 2002). Our results also showed that DNA loss in hg19 and mm10 in the hg19 genomic background was positively associated with genomic recombination. This is consistent with previous analyses that have identified an association between DNA loss and recombination (Nam and Ellegren 2012). Interestingly, we did not observe any association with recombination in the mm10 genomic background. This may be due to the decreased resolution used to calculate recombination rates and identify recombination hotspots in mouse compared to human (Brunschwig et al. 2012; International HapMap Consortium et al. 2007). For DNA gain hotspots we found that their associations with genomic features was less consistent across both species than DNA loss hotspots. For sources of DNA gain, mm10 and hg19 DNA gains were both positively associated with lineage-specific L1s. However, while lineage-specific SINEs were associated with hg19 DNA gain, in mm10 they were associated with DNA loss. This paradoxical finding is likely caused by two separate contributing factors. The first is that lineage-specific SINEs in mouse are not a major contributor to DNA gain compared to human, as their overall coverage levels are much lower (Chinwalla et al. 2002). The second is that lineage-specific SINEs accumulate in gene-rich open chromatin areas which also happen to strongly associate with DNA loss (Buckley et al. 2017). These differences in sources of DNA gain may explain divergence patterns in both species DNA gain dynamics; lineage-specific SINEs are associated with gene-rich/active genomic regions and lineage-specific L1s are associated with gene-poor silent regions such as LADs. Ultimately, this suggests that DNA is accumulating/turned over in different regions at different rates by otherwise conserved mechanisms of DNA gain. Collectively, our results show that DNA gain and loss is associated with specific genomic contexts, leading to differences in genome structure.

DNA gain and loss is non-random and may be a function of mammalian genome structure. However the evolutionary impact of DNA gain and loss is mainly determined by whether or not it affects particular phenotypes. To identify potentially impacted phenotypes we performed gene ontology (GO) enrichment analysis on genes in DNA gain and loss hotspots for biological process GO terms (Ashburner et al. 2000). Because we are interested in identifying whether DNA gain and loss may have driven lineage-specific divergence we compared the significance levels of GO term enrichment between our hotspot types. To do this we performed correlation analysis using the -log10 P-values for GO term enrichment as determined using a Fisher test combined with the ‘classic’ GO term enrichment algorithm (methods) (Alexa and Rahnenfuhrer 2016). Surprisingly our results showed the highest level of similarity between hg19 DNA gain and hg19 DNA loss (Fig. 6,S10). This is interesting because the overlap between hg19 gain and loss was not statistically significant (Fig. 4, S9). Moreover, when we compare hg19 DNA loss with mm10 DNA loss; gap annotations with a significant degree of overlap (Fig. 4, S9), we found that GO terms were not as similar, particularly in the mm10 genomic background (Fig. S10). Alternatively, enriched GO terms found in mm10 DNA gain hotspots appeared distinct from GO terms enriched in other DNA gain and loss hotspots. These results echo our above findings from comparing hotspot overlap, where mm10 gains were least likely to significantly overlap other hotspot types (Fig. 4,S9).

**Figure 6.**
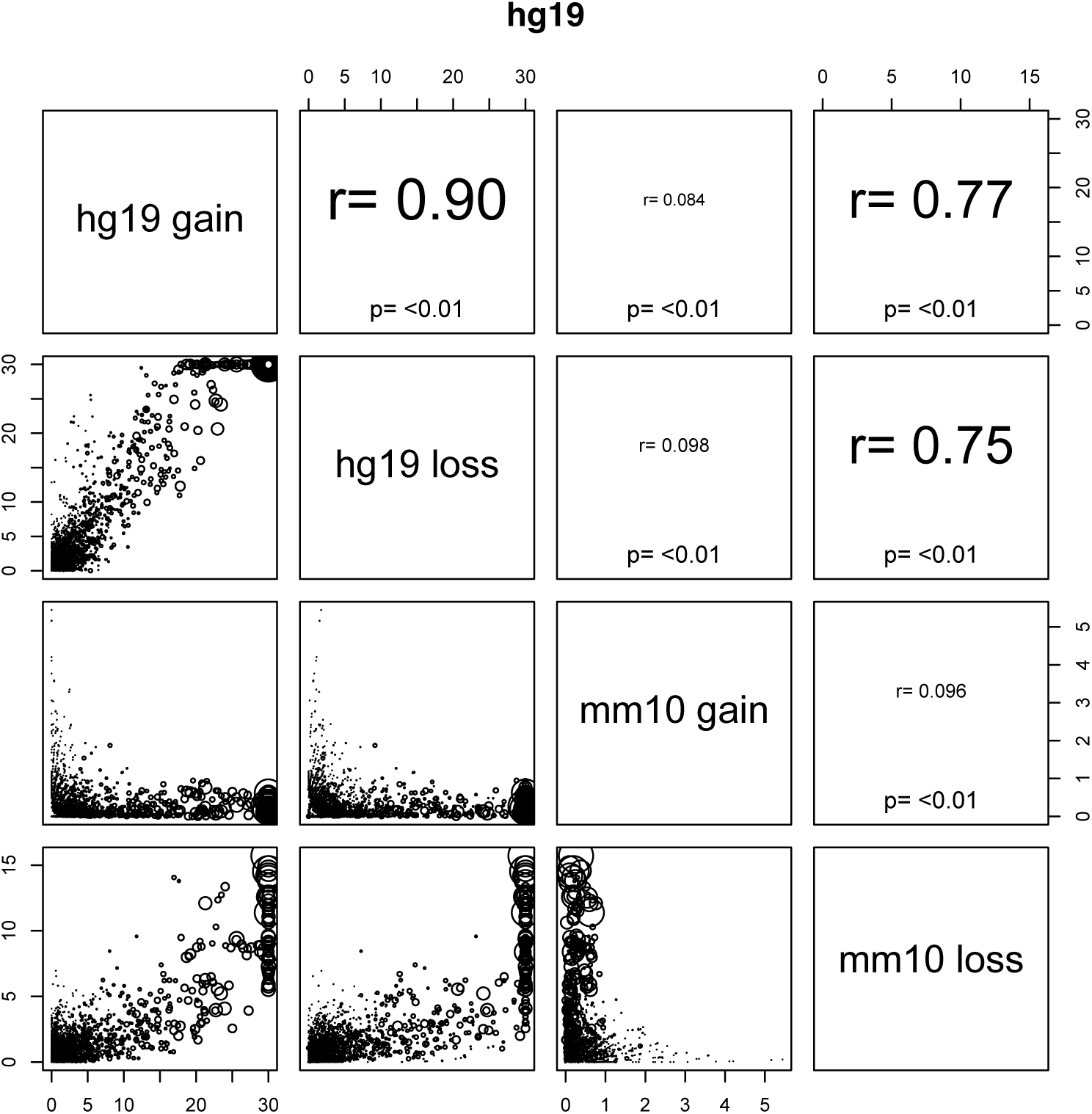
Over representation of biological process GO terms in gain and loss hotspots in hg19. The axes are marked according to -log10 P-values. The size of points represents the total number of annotations for each GO term.

To confirm our findings and examine the GO terms themselves, we calculated the proportion of significant terms that were descendants (child terms) of a high-order parent term. Child terms were identified as statistically significant at a FDR < 0.05 based on a Fisher test using the classic algorithm. Additionally, we extracted the 10 highest ranked terms discovered using the Fisher test combined with 3 other algorithms designed to reduce false positives generated by the inheritance problem (described in methods) (Table S3-S6) (Alexa et al. 2006; Grossmann et al. 2007). Statistically significant terms for hg19 gain and loss mostly belonged to cellular processes, metabolic processes, single organism processes and biological regulation (Fig. 7). For mm10, DNA loss hotspots were enriched for similar terms, including developmental processes, which were particularly enriched in the mm10 genomic background (Fig S11). However, mm10 gain in the hg19 background was only enriched for a single term and in the mm10 background mm10 gain was not enriched for any terms. The difference in these results is consistent with how DNA gain and loss events in human and mouse associate with regions of varying gene density and biological activity (Fig. 5).

**Figure 5.**
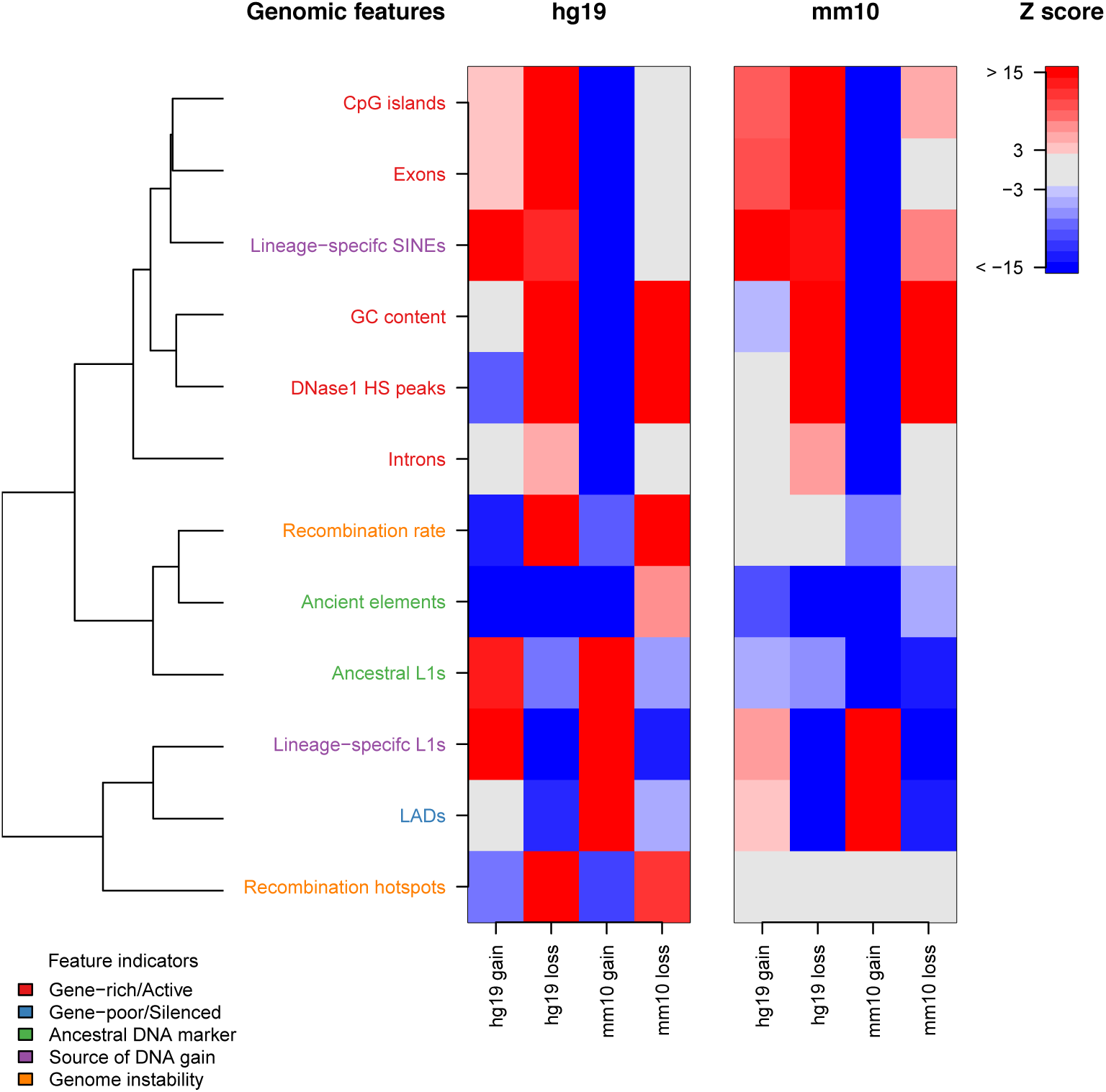
Association between genomic features and DNA gain or loss. Z scores are calculated using background distribution generated from 10000 permutations (methods). A positive association indicates that a particular gap annotation and genomic feature co-locate. Alternatively, a negative association indicates that the gap annotation and genomic feature occupy distinct genomic regions. DNaseI HS peaks (ENCODE Project Consortium et al. 2012), recombination hotspots (International HapMap Consortium et al. 2007; Brunschwig et al. 2012), LADs (Guelen et al. 2008; Peric-Hupkes et al. 2010), CpG islands (Tyner et al. 2016), gene annotations (Carlson 2015, 2016) and Retrotransposons (Smit et al. 2015) were measured in each as coverage per 200 kb. Recombination rates were measured as the mean bin-wise recombination rate (International HapMap Consortium et al. 2007; Brunschwig et al. 2012). GC content was measured as the proportion of G or C nucleotide residues in chain-blocks per bin (Team TBD 2014a,b). Genomic features are classified intro groups of feature indicators based on distinct aspects of genome biology they are known to associate with. The dendrogram represents spatial clustering of genomic features across both genomes,where two tightly clustered genomic features in the dendrogram are genomic features that tend to be co-located. The dendrogram was generated from a correlation matrix that consisted of pair-wise correlations between each feature across both binned genomes.

**Figure 7.**
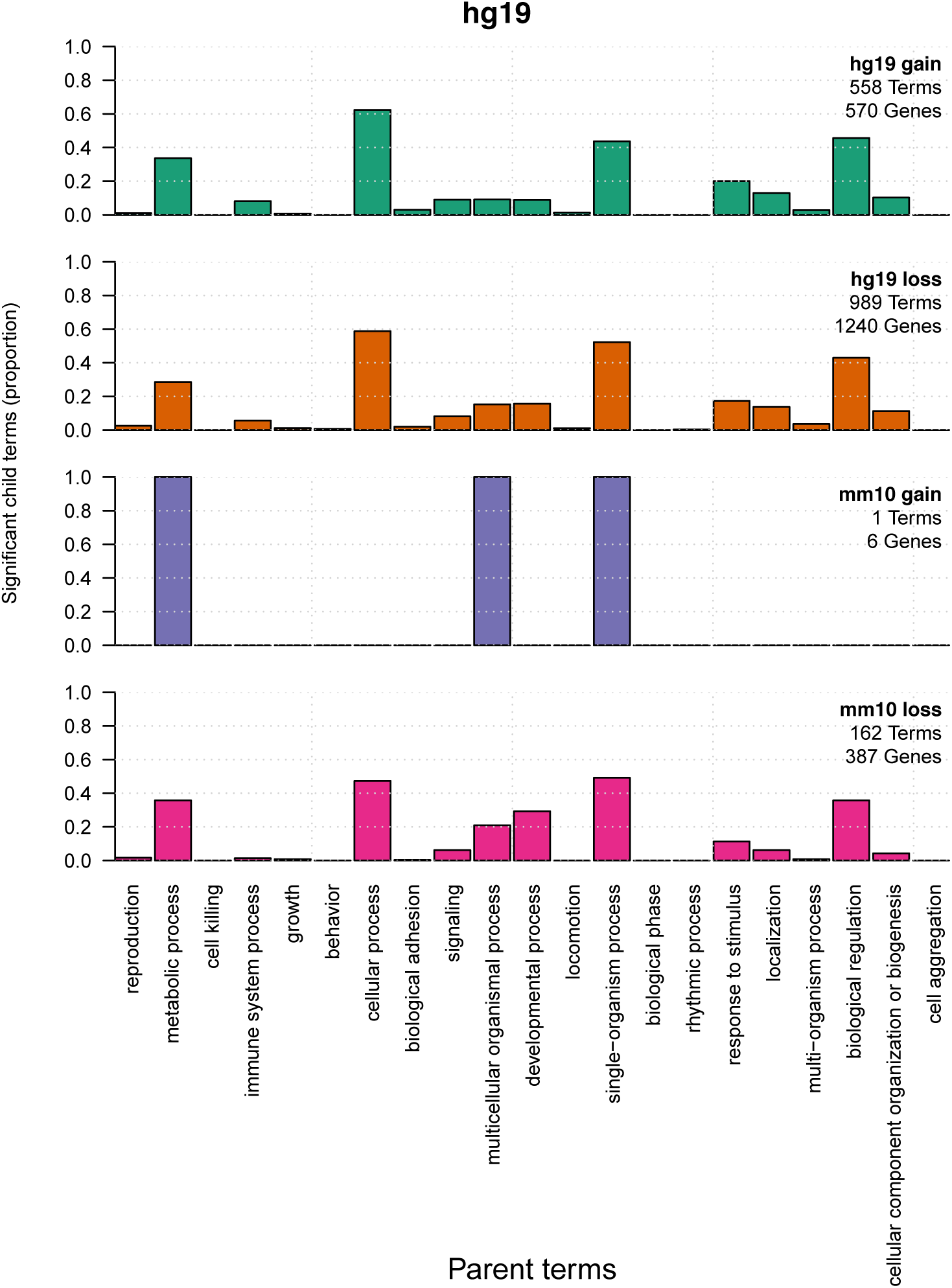
Significant biological process GO terms in hg19 background. Parent terms were the top level biological process GO terms while child terms were those beneath each parent term. Child terms were identified as significant at a FDR < 0.05 based on a Fisher test using the ‘classic’ algorithm. The Y axis represents the proportion of child GO terms that belong to each parant GO term. Proportions don’t add up to 1 because some child GO terms are shared between parent GO terms. We have also shown the number of non-redundant GO terms and genes annotated with significant GO terms for each gap annotation.

Interestingly, while the genomic distributions of each hotspot type differed, their associated significant GO terms were highly similar. This may be caused by genes that contribute to similar biological processes being tightly clustered and located within regions that consist of overlapping hotspot types. To determine if this was the case we compared non-redundant statistically significant child terms and gene annotations across each hotspot type (Fig S12). We found that the vast majority of genes annotated with significant GO terms were unique to a particular hotspot type. In contrast to this, the GO terms were more likely to be shared across hotspot types. This suggests that DNA gain and loss tend to associate with different genes that contribute to the same biological processes. Together our results show that particular biological processes are either prone to DNA gain or loss or are instead highly robust and able to withstand high levels of genomic turnover.

## Discussion

### Genome-wide DNA gain and loss dynamics

Estimating the total amount of DNA turnover across two separate lineages over a time span of approximately 90 million years is a challenging task (Hedges et al. 2006). After this divergence period as little as 40% of the extant human genome shares ancestry with mouse, suggesting that at least 60% has been turned over in either lineage. In order to understand gain and loss dynamics we must be able to correctly assign this non-aligning portion of the human genome as either human gain or mouse loss. Chinwalla et al. (2002) and Hardison et al. (2003) used an approach similar to our recent transposon based method. They used a set of lineage-specific transposons in human and mouse to identify regions of DNA gain. From this, the remaining non-aligning portion of one genome was assumed to be lost from the other. To confirm this approach, Chinwalla et al. (2002) checked to see if their inferred genome-wide rates of DNA loss were consistent with local estimates. They used the following equation;

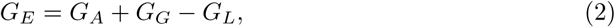

where *G*_*E*_ is the size of the extant genome, *G*_*A*_ is the size of the ancestral genome, *G*_*G*_ is the amount of lineage-specific genome gain and *G*_*L*_ is the amount of lineage-specific genome loss. For human and mouse they solved the equation for *G*_*L*_ where they estimated ancestral genome size within a range similar to the extant human genome size. This was chosen because it was similar to the average genome size for mammalian outgroup species. Estimates showed that DNA loss in mouse was almost double that of human, and consistent with the difference in the number of non-aligning non-recent transposon bases in each genome. While these estimates were consistent with expectations based on the assumption that non-aligning non-recent transposon regions were ancestral, their ancestral state remained unverified. Conversely, our ancestral based approach aimed to directly verify the ancestry status of non-aligning regions between human and mouse. This was achieved by using a wide variety of outgroup species alignments not available to Chinwalla et al. (2002) and Hardison et al. (2003) at the time of their analysis. In human, our results revealed that indeed many of the non-aligning non-recent transposon bases overlapped ancestral elements. However, approximately 168 Mb remained ambiguous (Table 2) which was more than double the 5.8% of the total non-aligning human genome, the fraction of known ancestral bases not supported by ancestral elements (Table 1). As stated in the results, this discrepancy was most likely caused by incorrect identification of DNA gain events or misidentification of ancestral elements. It is important to realise that the ancestral element based approach has its limits, as orthologous sequences between species have the potential to diverge beyond recognition. This was the most likely reason that ancestral element detection in mouse was so much lower than in human, as the genome-wide substitution rate in mouse is approximately twice that of human.

An alternative way to verify the recent transposon based method was to use our estimated DNA loss rates to solve for *G*_*A*_ and to compare this to other estimates of ancestral genome sizes. After the mouse genome was completed many other mammalian genome projects also reached completion, allowing for the development of ancestral genome reconstruction techniques. While ancestral genome reconstruction is based on alignment it is much less susceptible to errors than our detection of ancestral elements. Instead of performing alignments directly between human or mouse and each individual outgroup species, it uses alignments between groups of more closely related species to build a phylogeny of ancestral states (Blanchette et al. 2004; Ma et al. 2006). Recently, Kim et al. (2017) estimated an ancestral euarchontoglires genome of 2.67 Gb in an analysis involving 19 placental mammals. Using equation 2 and solving for *G*_*A*_ with extant genome sizes from Table 1 and gain and loss rates calculated by the recent transposon method (Table 2), we get estimated ancestral genome sizes of 2.64 Gb and 2.66 Gb for human and mouse respectively. Together our findings in the context of various other methods support the use of recent transposons to analyse DNA gain and loss dynamics.

While the recent transposon method provides an accurate estimate of DNA gain and loss dynamics it is important to realise these estimates are only a lower bound on the the total amount of DNA turnover since divergence. This is because both our analysis and previous analyses relied heavily on the assumption of parsimonious genome evolution, where lineage-specific gain and loss patterns are based on the fewest possible evolutionary changes. Unfortunately, in our case the assumption of parsimonious genome evolution is likely to cause various events to be hidden. For example, if a particular region underwent lineage-specific DNA gain that was subsequently lost, both the gain and loss events will not be detected. Additionally, DNA loss occurring in both lineages at the same loci would also go undetected. Depending on the frequency and magnitude of the above events we have likely underestimated the total amount of DNA gain and loss. A possible way to overcome this problem is to adopt model based approaches similar to those used in phylogenetic analyses. These approaches use a substitution model along with maximum likelihoods or Bayesian inference to allow for varying rates of evolution across lineages and sites (Yang and Rannala 2012). However, given our current lack of understanding of the non-coding portion of the genome such an approach for estimating DNA turnover is likely to yield highly questionable results.

### Evolutionary impact of large scale DNA gain and loss

During genome evolution the spectrum of possible mutations is extremely broad, ranging from single nucleotide substitutions all the way up to Mb-sized rearrangements and translocations. Importantly, the genomic distribution of events at each level of the mutation spectrum is non-random and highly context-dependent. Moreover, the regional susceptibility and tolerance to a particular mutation type is a mixture of various genomic and epigenomic features and selective pressures (Makova and Hardison 2015). To understand the evolutionary impacts and trajectories of DNA gain and loss dynamics we analysed their genomic distributions in the context of various genomic features and biological processes.

In mammals synteny is highly conserved due to the frequent reuse of chromosome rear-rangement breakpoints throughout their evolution (Murphy et al. 2005). Since chromosome rearrangement breakpoints were located outside of nets, many DNA gain and loss events went undetected (S1-S2). Instead, we most likely identified regions where gain and loss dynamics impacted on local architecture, such as the genomic distances between neighbouring genes or intron size. However, due to the difficulty in mapping DNA gain and loss events across large evolutionary time scales, the impact of DNA gain and loss at this scale remains largely unknown. Our strategy has therefore allowed us for the first time to measure regional variation in DNA gain and loss across genome structures that have been resistant to large structural rearrangements. Our results revealed that DNA gains and losses in human and mouse were associated with the same kinds of features; DNA gains were most associated with L1 accumulation in gene poor regions with low biological activity while DNA losses occurred mostly in highly active gene-rich regions. Previous analyses have shown that genome organisation between human and mouse is largely conserved, where lineage-specific L1s and SINEs tend to accumulate in similar regions in different species (Buckley et al. 2017). Our results suggest that rather than certain types of events driving genome divergence, it is instead the rate at which each particular event type occurs that drives divergence. For example, mouse has a much higher deletion rate than human and a larger number of active L1s. This would suggest that particular regions in the mouse are growing or shrinking much more than in the human genome while their sequence composition remains similar. Alternatively, DNA gain rates were especially enriched on the X chromosome in both species with some degree of regional overlap (Fig. 4,S9). This is consistent with the high concentration of L1s that play a role in X inactivation (Chow et al. 2010).

Despite the amount of structural divergence between human and mouse, it is difficult to identify how much impact this might have on evolution at the level of phenotype. Interestingly, Human DNA gains and losses and mouse DNA losses all occurred near genes involved in fundamental cellular/metabolic processes. Because cellular/metabolic process genes likely evolved earlier in animals and probably have house keeping functions, their regulation is also likely highly conserved (Lowe et al. 2011). This suggests that for the most part the accumulation of DNA gains and losses have had little impact on phenotypic change. However, for some mouse DNA losses the case may be different, as in the mm10 genomic background they mostly occurred near genes involved in developmental processes. Developmental processes may be linked to traits that could have potentially undergone divergence, such as mouse-specific morphological characteristics. While this is an attractive idea, an analysis of regulatory element evolution shows that lineage-specific regulatory innovation for development occurred prior to human and mouse divergence (Lowe et al. 2011). Therefore, throughout mammalian evolution regulatory elements for development and cellular processes have likely remained intact while nearby DNA has been frequently turned over. Ultimately, given that we are able to detect little phenotypic impact where there are vast amounts of DNA turnover, our findings raise questions regarding the proportion of the human genome that is under selection and indeed ‘functional’.

Topological associated domains (TADs) are a particular aspect of genome-organisation that may be affected by our detected DNA gains and losses. TADs are Mb-sized units of genome organisation that consist of highly self-interacting DNA. For example, two distant loci within a single TAD are much more likely to interact with each other than two loci that are near each other but happen to be located within different TADs (Dixon et al. 2012). Because TAD boundaries associate with other domain boundaries linked to gene regulation, such as LADs, they are often considered as distinct autonomously regulated regions (Sexton and Cavalli 2015). Since TADs are organised along a linear stretch of DNA, it is possible that their organisation is somewhat dependent on genomic distances between co-regulated features. This suggests that increased lineage-specific DNA gain and loss may cause TAD structures to diverge. One way this could happen is by removing TAD boundaries through deletion, which would subsequently cause adjacent TADs to merge (Hnisz et al. 2016). Alternatively, increases in the genomic distance between the edges of a single TAD could potentially promote the formation of a new boundary. These scenarios are more likely to have occurred in mouse rather than human, where DNA gain and loss in mouse is much more regionally clustered, ultimately causing larger deviations from regional gain and loss equilibrium. In vertebrates, *Hox* clusters are located between two adjacent TADs that most likely diverged from a single TAD leading to the evolution of the vertebrate *Hox* bipartite regulatory system (Acemel et al. 2016). This new TAD structure has made it possible for *Hox* genes to receive new inputs from distal enhancers contributing to the evolution of limb development and anteroposterior axis pattering (Lonfat and Duboule 2015). So while regulatory innovation at the level of individual elements may have slowed prior to human and mouse divergence, changes in TAD structure may cause ancestral enhancer elements to be co-opted in developmental processes driving lineage-specific phenotypic evolution.

## Conclusion

There are four key points from our results. First, hot spots for DNA gains and losses occur in different compartments; loss hot spots in open chromatin/regulatory regions and gain hot spots in heterochromatin. Because DNA loss is caused by repair of DNA Double Stranded Breaks (DSB) (Gasior et al. 2006), this means that L1 ORF2p activity can both cause DNA gains and losses as a cause of DSB. However, this does not mean that gains and losses do not occur in the same regions. Second, mouse SINEs are strongly associated with DNA loss, indicating that losses in regulatory regions are accompanied by SINE insertions suggesting that there is extensive “churning” or turnover of sequences in these regions. The observed differences in associations between lineage-specific SINEs and gain and loss in mouse and human are likely due to differential expansion of LINEs vs SINEs in the two lineages. Thus, regional/species specific variation in DNA gain and loss are primarily driven by clade specific/recent transposons interacting with open chromatin either in the male germ line, female germ line or early embryo. Third, the X chromosome is largely devoid of loss hot spots, but has many gain hot spots, consistent with a continuing selection for insertion of L1 elements required for X inactivation. Fourth, the observed autosomal divergence of gain and loss hot spot patterns in proximity to genes supports a model in which selection of altered developmental/regulatory mechanisms (based on GO term results) occurs as a result of transposon driven DNA gain and loss. This has implications for our views regarding the “functional” proportion of the genome that is under selection and contributing to phenotypic divergence.

## Additional Files

Additional file 1 — Supplementary information

## Author’s contributions

R.M.B., R.D.K., and D.L.A. designed research; R.M.B. performed research; and R.M.B., R.D.K., and D.L.A. wrote the paper.

## Acknowledgements

We would like to thank Steve Pederson, Rick Tearle, Jonathan Henry Jacobsen, Lu Zeng and Zhipeng Qu for their helpful discussion throughout the research process and Catisha Coburn for help with editing the manuscript.

## Availability of data and materials

